# VALIS: Virtual Alignment of pathoLogy Image Series

**DOI:** 10.1101/2021.11.09.467917

**Authors:** Chandler D. Gatenbee, Ann-Marie Baker, Sandhya Prabhakaran, Robbert J. C. Slebos, Gunjan Mandal, Eoghan Mulholland, Simon Leedham, Jose R. Conejo-Garcia, Christine H. Chung, Mark Robertson-Tessi, Trevor A. Graham, Alexander R.A. Anderson

## Abstract

Spatial analyses can reveal important interactions between and among cells and their microenvironment. However, most existing staining methods are limited to a handful of markers per slice, thereby limiting the number of interactions that can be studied. This limitation is frequently overcome by registering multiple images to create a single composite image containing many markers. While there are several existing image registration methods for whole slide images (WSI), most have specific use cases. Here, we present the Virtual Alignment of pathoLogy Image Series (VALIS), a fully automated pipeline that opens, registers (rigid and/or non-rigid), and saves aligned slides in the ome.tiff format. VALIS has been tested with 273 immunohistochemistry (IHC) samples and 340 immunofluorescence (IF) samples, each of which contained between 2-69 images per sample. The registered WSI tend to have low error and are completed within a matter of minutes. In addition to registering slides, VALIS can also using the registration parameters to warp point data, such as cell centroids previously determined via cell segmentation and phenotyping. VALIS is written in Python and requires only few lines of code for execution. VALIS therefore provides a free, opensource, flexible, and simple pipeline for rigid and non-rigid registration of IF and/or IHC that can facilitate spatial analyses of WSI from novel and existing datasets.

## Introduction

Cellular interactions and the structure of the tumor microenvironment can affect tumor growth dynamics and response to treatment (Gallaher, Enriquez-Navas, Luddy, Gatenby, & Anderson, 2018; Heindl et al., 2018; Lewis et al., 2021). Interactions and the effect of tissue structure can be elucidated via spatial analyses of tumor biopsies, although there are many challenges. Among these is the limited number of markers that can be detected on a single tissue section. This can be overcome by repeated cycles of staining on the same tissue section, or by staining serial slices for different subsets of markers. However, the captured images will likely not align spatially, due to variance in tissue placement on the slide, tissue stretching/tearing/folding, and changes in physical structure from one slice to the next. Without accurate alignment, spatial analyses remain limited to the number of markers that can be detected on a single section. While there are methods that can stain for a large number of markers on a single slide, they are often highly expensive, destructive, and require considerable technical expertise (Angelo et al., 2014; Gerdes et al., 2013; Giesen et al., 2014; Goltsev et al., 2018; Hennig, Adams, & Hansen, 2009). Furthermore, much archival tissue is available that has limited stains per slice.

Image registration is the process of aligning one image to another such that they share the same coordinate system, and therefore offers the potential to align histology images. However, a pre-requisite for successful image registration is that the images look similar, but this requirement is rarely satisfied in histological images. The reasons for this may include: spatial variation in color intensity due to markers binding in different regions of the slide; lack of a common marker across images (in the case of IHC); inter-user or inter-platform variation in staining intensity; tissue deformations (e.g. stretching, folds, tears); unknown order of serial sections; large numbers of images; and massive file sizes, often several GB when uncompressed (Figure 1).

**Figure 1.**
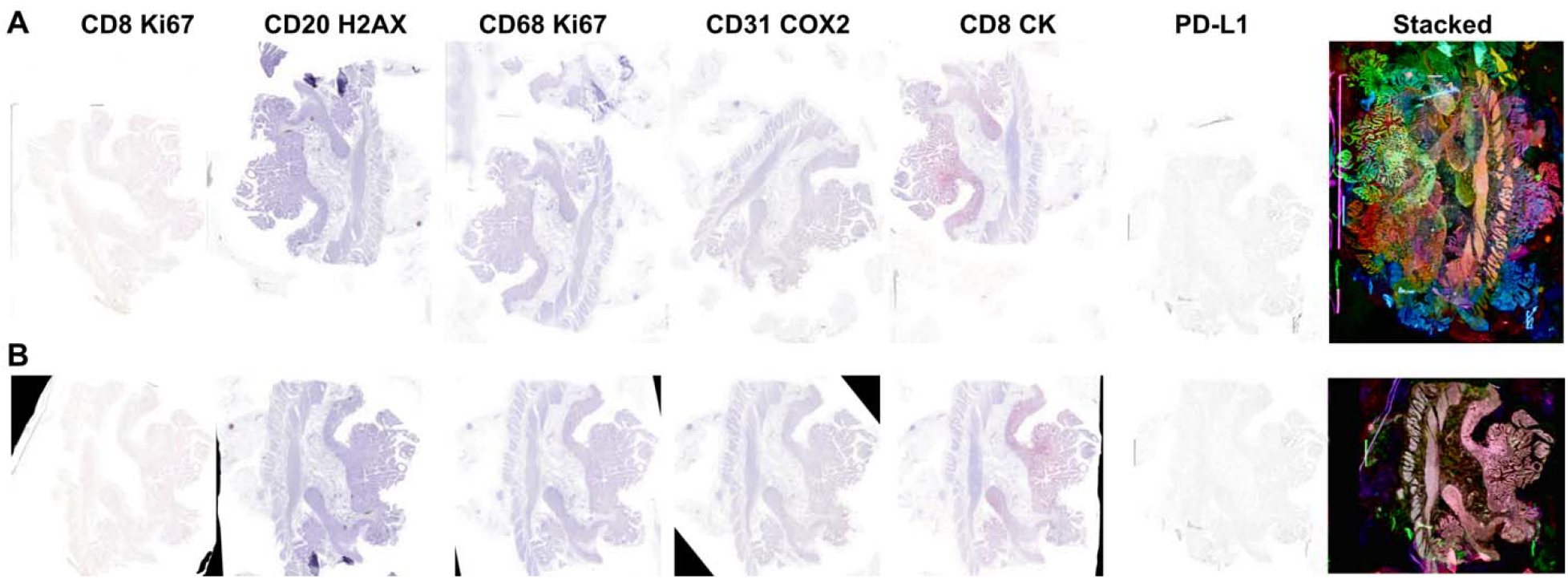
Example of a challenging dataset successfully registered by VALIS. VALIS handles potential batch effects from IHC images that would otherwise make image registration challenging. Such batch effects include large displacements (rotation, translation, etc.); deformations (stretches, tears); and spatial variation in color and luminosity due to differing spatial distributions of markers and/or different staining protocols. Large file sizes also present challenges to registering whole slide images (WSI). **A**) Six serial slices of a colorectal adenoma were stained by three different individuals, with each marker stained with Fast Red or DAB. Note the substantial spatial variation in color and brightness, due to the heterogeneous spatial distribution of different cell types (each type stained with a different marker), and different staining protocols where some images are heavily stained and others lightly stained. The rightmost image shows the result of stacking the un-registered images, where each color shows the normalized inverted luminosity of each image. Each slide is also too large to open in memory, with each being ∼32GB when uncompressed. **B**) Left: Alignment of the same slides using VALIS. Right: Image stack after image registration using VALIS. The transformations found by VALIS can subsequently be used warp each of the 32Gb slides, which can be saved as ome.tiff images for downstream analyses.

There are several existing methods to register histological images, many of which have been reviewed in (Jiří Borovec, Muñoz-Barrutia, & Kybic, 2018; Paknezhad et al., 2020). Some are limited to hematoxylin and eosin (H&E) staining (Arganda-Carreras et al., 2006; du Bois d’Aische et al., 2005; Kiemen et al., 2020; Wang, Ka, & Chen, 2014), while others are designed to work with slides stained for different markers (J. Borovec, Kybic, Bušta, Ortiz-de-Solórzano, & Muñoz-Barrutia, 2013; Deniz, Toomey, Conway, & Bueno, 2015; Kybic & Borovec, 2014; Kybic, Dolejší, & Borovec, 2015; Obando et al., 2017; Song, Treanor, Bulpitt, & Magee, 2013). Some are designed to align only 2 slides (Levy, Jackson, Haudenschild, Christensen, & Vaickus, 2020), while others can align multiple slides (Kiemen et al., 2020; Paknezhad et al., 2020). There also exist several methods to register immunofluorescence (IF) images, which can be an easier task as each image usually contains a DAPI channel that stains for nuclei (Muhlich, Chen, Russell, & Sorger, 2021). Most seem to require the user be able open the slides, apply the transformation, and then save the registered image. All these methods also focus exclusively on IHC or IF images. Thus, while each method has many benefits, they also have limitations that can reduce their use cases.

Here, we present the Virtual Alignment of pathoLogy Image Series (VALIS), which aims to combine the best of current approaches while remaining easy to use. VALIS provides the following 7 advantages: 1) The tool is flexible and unique, as it is able to align both immunohistochemistry (IHC) and immunofluorescence (IF) images, whole slide images (WSIs) or regions of interest (ROIs), H&E images or collections of different markers, serial slices and/or cyclically stained images (tested using 11-20 rounds of staining); 2) It can register any number of images, find rigid and/or non-rigid transformations, and apply them to slides saved using a wide variety of slide formats (via Bio-Formats or OpenSlide), and then save the registered slides in the ome.tiff format (Gohlke, 2021; Goldberg et al., 2005; Goode, Gilbert, Harkes, Jukic, & Satyanarayanan, 2013; Linkert et al., 2010; Martinez & Cupitt, 2005); 3) VALIS is designed to be extendable, giving the user the ability to provide additional rigid and/or non-rigid registration methods; 4) The user may also provide transformations found using other methods but still use VALIS to warp and save the slides; 5) The transformations found by (or provided to) VALIS can be applied not only to the original slide, but also processed versions of the slide (e.g. ones that have undergone stain segmentation) which could be merged; 6) The transformations found by VALIS can also be used to warp point data, such as cell positions (Figure 5); 7) VALIS is designed to be easy to use, requiring only a few lines of code in Python, or a few clicks of a mouse in a user-friendly graphical user interface (GUI). Thus, VALIS provides a simple and fully automated registration pipeline to open, register, and save a series of pathological images.

## Results

## Method Overview

### Reading the slides

Whole slide images (WSI) can be challenging to work with because they are saved using a wide variety of formats. They are often very large in size (greater than 70,000 pixels in width and height) and have multiple channels. The resulting uncompressed file is frequently on the order of 20GB in size, which precludes opening the entire image directly. To address the issues of working with WSI, VALIS uses Bio-Formats and OpenSlide (Goode et al., 2013; Linkert et al., 2010) to read each slide in small tiles, covert those tiles to *libvips* images, and then combine the tiles to rebuild the entire image as single whole-slide *libvips* image (Linkert et al., 2010; Martinez & Cupitt, 2005). As *libvips* uses “lazy evaluation”, the WSI can then be warped without having to load it into memory, making it ideal for large images. Using this approach, VALIS is thus able to read and warp any slide that Bio-Formats can open.

### Preprocessing

VALIS uses tissue features to find the transformation parameters, and therefore a lower resolution version of the image is used for feature detection and finding the displacement fields used in non-rigid registration. The lower resolution image is usually acquired by accessing an upper level of an image pyramid. However, if such a pyramid is unavailable, VALIS can use libvips to rescale the WSI to a smaller size. The images used for feature detection are usually between 500-2000 pixels in width and height. Prior to feature detection, all processed images are re-sized such that all have the same largest dimension (i.e. width or height).

For image registration to be successful, images need to look as similar as possible. In the case of IF, the DAPI channel is often the best option to use for registration. However, unless one is only working with H&E, a preprocessing method to make IHC images look similar must be used. The default method in VALIS is to first to re-color each image to have the same hue and colorfulness. This is accomplished by converting the RGB image to the polar CAM16-UCS colorspace (C. Li et al., 2017), setting C=100 and H=0 (other values can be used), and then converting back to RGB. The transformed RGB image is then converted to greyscale and inverted, such that the background is black, and the tissue bright. After all images have been processed (IHC and/or IF), they are then normalized such that they have similar distributions of pixel values. The normalization method is inspired by (Khan, Rajpoot, Treanor, & Magee, 2014), where first the 5^th^ percentile, average, and 95^th^ percentile of all pixel values is determined. These target values are then used as knots in cubic interpolation, and then the pixel values of each image are fit to the target values.

### Rigid Registration

VALIS provides a novel pipeline to rigidly align a large number of unsorted images, using feature detecting and matching. A major benefit of using feature-based registration is that it can cope with large displacements, and thus does not require the images to already be somewhat aligned. The default feature detector and descriptor are BRISK and VGG, respectively (L. Li, Huang, Gu, & Tian, 2003; Simonyan, Vedaldi, & Zisserman, 2014). Features can then be matched using brute force, with outliers removed using the RANSAC method (Fischler & Bolles, 1981). The remaining “good” matches can then be used to find the rigid transformation parameters.

#### Ordering images

If the order of images is unknown, the following steps are used to determine the order in which images will be sequentially aligned. First, feature matching and filtering is performed for each pair of images. Next, the feature matches are used to construct a similarity matrix **S**, where the default similarity metric is simply the number of good matches between each pair of images. **S** is then standardized such the maximum similarity is 1, creating the matrix **S**^**′**^, and then converted to the distance matrix, **D**=1− **S**^**′**^. Hierarchical clustering is then performed on ***D***, generating a dendrogram *T*. The order of images can then be inferred by optimally ordering the leaves of *T*, such that most similar images are adjacent to one another in the series (Bar-Joseph, Gifford, & Jaakkola, 2001). This step can be skipped if the order of images is known.

#### Warping images

Once the order of images has been determined, VALIS finds the transformation matrices that will rigidly warp each image to the previous image in the series. That is, each image *I*_*i*_ will have an associated transformation matrix *M*_*i*_ that rigidly aligns it to image *I*_*i*−*1*_, where *i* is the position of the image in the series. While RANSAC does an excellent job of removing poor matches, those mismatched features are sometimes considered inliers and thus potentially used to estimate transformation matrices. Including the coordinates of such mismatched features will produce poor estimates of the transformation matrices that will align feature coordinates. To avoid this, only features that are found in the image and its neighbors are used. That is, the features used to align image *I*_*i*_ and *I*_*i*−*1*_ are the features that *I*_*i*_ also has in common with *I*_*i+1*_ and thus consequently that *I*_*i*−*1*_ also has in common with *I*_*i+1*_ The assumption here is that features which are found in *I*_*i*_ and its neighborhood are shared because they are strong tissue landmarks, and thus ideal for alignment. This approach may be thought of as using a sliding window to filter out poor matches by using only features shared within an image’s neighborhood/community. The coordinates of the filtered matches are then used to find the transformation matrix (*M*_*i*_) that rigidly aligns *I*_*i*_ to *I*_*i*−*1*_.

#### Final optional alignment optimization

After warping all images using their respective rigid transformation matrices, the series of images has been registered. However, one can optionally use an intensity-based method to improve the alignment between *I*_*i*_ and *I*_*i*−*1*_. The default in VALIS is to maximize Mattes mutual information between the images, while also minimizing the distance between matched features (Lowekamp, Chen, Ibanez, & Blezek, 2013). Once optimization is complete, *M*_*i*_ will be updated to be the matrix found in this optional step. This step is optional because the improvement (if any) may be marginal (distance between features improving by fractions of a pixel), and it is time consuming.

#### Non-Rigid Registration

Non-rigid registration involves finding 2D displacement fields, *X* and *Y*, that warp a “moving” image to align with a “fixed” image by optimizing a metric. As the displacement fields are non-uniform, they can warp the image such that local features align better than they would with non-rigid transformations (Crum, Hartkens, & Hill, 2004). However, these methods require that the images provided are already somewhat aligned. Therefore, once VALIS has rigidly registered the images, they can be passed on to one of these non-rigid methods. VALIS can conduct this non-rigid registration using one of three methods: Deep Flow, SimpleElastix, or Groupwise SimpleElastix (Klein, Staring, Murphy, Viergever, & Pluim, 2010; Marstal, Berendsen, Staring, & Klein, 2016; Shamonin et al., 2013; Weinzaepfel, Revaud, Harchaoui, & Schmid, 2013). In the case of the first two methods, images are aligned towards the image at the center of the series. For example, given N images, the center image is 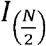. Therefore, 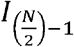 is aligned to 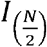, then 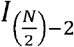 is aligned to the non-rigid warped version of 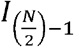, and so on. Each image’s displacement fields, *Xi* and *Yi*, are built through composition. For the third method (Groupwise SimpleElastix), this process of aligning pairs of images and composing displacement fields is not necessary, as it uses a 3D free-form B-spline deformation model to simultaneously register all the images.

#### Warping and Saving

Once the transformation parameters *M*_*i*_, *X*_*i*_, and *Y*_*i*_ have been found, they can be scaled and used to warp the full resolution image, which is accomplished using *libvips*. The warped full resolution image can then be saved as an ome.tiff image, with the ome-xml metadata being generated by *tiffile* and saving done using *libvips* (Gohlke, 2021; Goldberg et al., 2005; Linkert et al., 2010). Once saved as an ome.tiff, the registered images can be opened and analyzed using open-source software such as QuPath (Bankhead et al., 2017) or commercially available software, such as Indica Labs HALO® (Albuquerque, NM, USA) image analysis software. As the ome.tiff slides can be opened using *libvips* or Bio-Formats, one can also use the aligned slide in a more tailored analysis using custom code.

### Registration Validation

To test the robustness of VALIS, we performed image registration on 613 samples, with images capture under a wide variety of conditions (Figures 2). Each sample had between 2 to 69 images; 273 were stained using immunohistochemistry (IHC), and 340 using immunofluorescence (IF); 333 were regions of interest (ROI) or cores from tumor microarrays (TMA), while 280 were whole slide images (WSI); the original image dimensions ranged from 2656 × 2656, to 104568 × 234042 pixels in width and height; 162 underwent stain/wash cycles, 451 were serial slices; 49 came from breast tumors, 109 from colorectal tumors, 156 from squamous cell carcinoma of the head and neck (HNSCC), and 299 from ovarian tumors.

**Figure 2.**
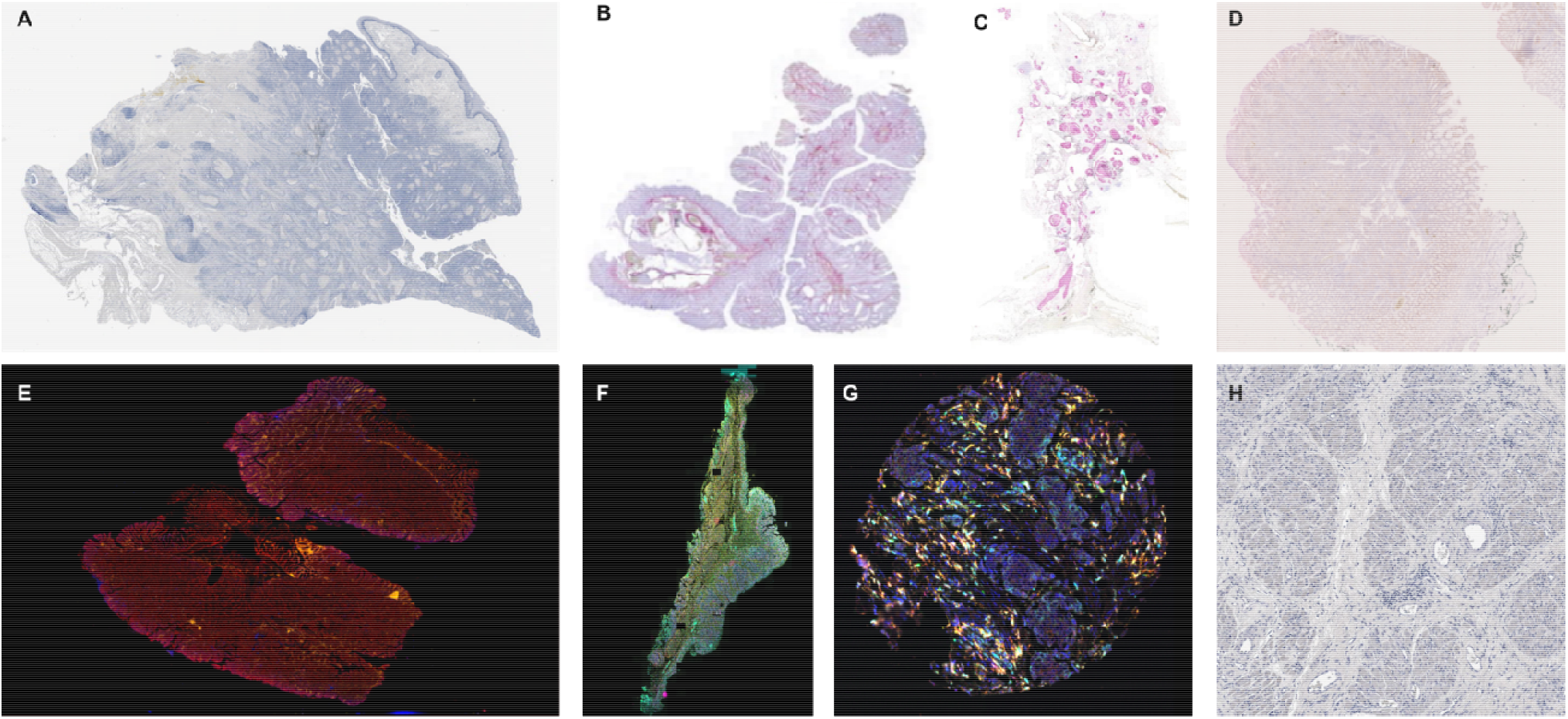
Example of images used to test VALIS. **A**) Squamous cell carcinoma of the head and neck (HNSCC). This sample set included 4 marker panels, each of which included between 13-20 markers stained using IHC. A single slice underwent the corresponding number of stain wash cycles, but all 69 images collected from the 4 panels have also been co-registered. **B**) Human colorectal carcinoma or adenoma IHC serial slices, each with 1-2 markers per slide, and 6 slides per sample. **C**) DCIS and invasive breast cancer serial slices, 1-2 markers per slide (stained using IHC), 7 slides per sample. **D**) Human colorectal carcinomas and adenomas, stained using RNAscope, 1-2 markers per slide, 5 slides per sample. **E**) Human colorectal carcinomas and adenomas, stained using cyclic immunofluorescence (CyCIF), 11-12 images per sample. **F**) Human colorectal carcinomas and adenomas stained using immunofluorescence, 2 slides per sample. **G**) In addition to registering WSI, VALIS can also be used to register images with cellular resolution, such as cores from an immunofluorescent tumor microarray (TMA) taken from human ovarian cancers (2 slides per sample), or **H**) 40x regions of interest from HNSCC samples, taken from images in the same dataset in panel A.

**Figure 3.**
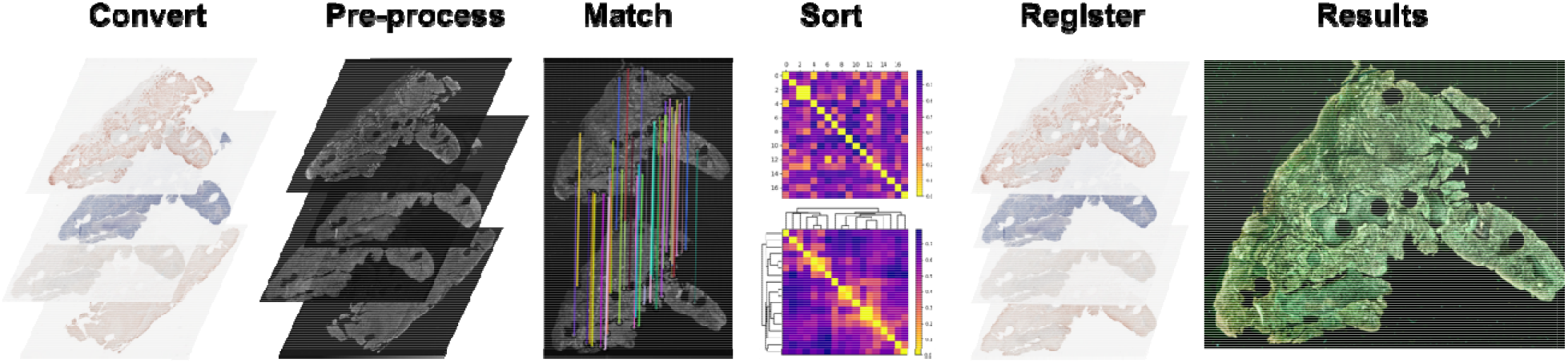
Overview of the VALIS alignment pipeline. VALIS uses Bio-Formats to read the slides and convert them to images for use in the pipeline. Once converted from slides, images are processed and normalized to look as similar as possible. Features are detected in each image and then matched between all possible pairwise combinations. Feature distances are used to construct a distance matrix, which is then clustered and sorted, ordering the images such that each image should be adjacent to its most similar image. Once ordered, the images are registered serially, first using rigid transformations, and then (optionally) with non-rigid transformations. The results of the registration can be viewed by overlaying the processed images. Once registration is complete, the slides can be warped and saved as ome.tiff images.

For each image, registration error was calculated as the median distance (μm) between the features in the image and the corresponding matched features in the previous image (see Methods section for more details) (Figure 4). The registration error of the sample was then calculated as the average of the images’ registration errors, weighted by the number of matched features per pair of images. The registrations provided by VALIS substantially improved the alignments between images, particularly in the case of serial IHC (Figure 4).

**Figure 4.**
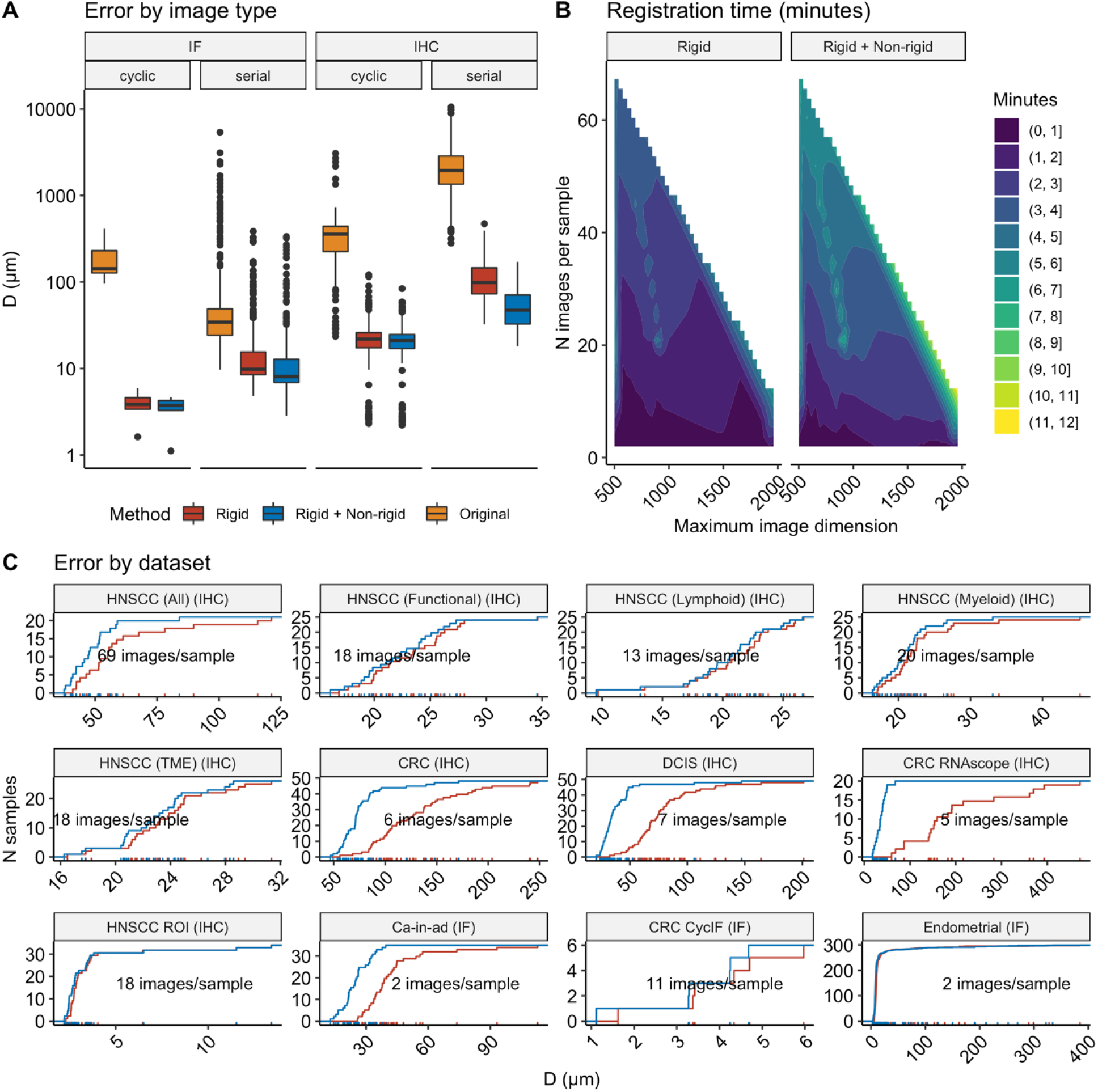
Results of registering images from the datasets shown in Figure 2, which were captured from a variety of tissues, protocols, imaging modalities, and resolutions. **A**) Boxplots showing the distance (μm) between matched features in the full resolution slides, before registration (yellow), after rigid registration (red), and then non-rigid registration (blue). **B**) Median amount of time (minutes) taken to complete registration, as a function of processed image’s size (by largest dimension, on the x-axis) and the number of images being registered. These timings include opening/converting slides, pre-processing, and intensity normalization. **C**) Empirical cumulative distribution plots of registration error for each image dataset.

## Applications

Alignment error was low for samples that underwent cyclic immunofluorescence (CyCIF), with an average distance between matched features (in the full resolution slide) being 2-6 μm apart. In these cases, the quality of the image registration was high enough that cell segmentation and phenotyping could be performed, as shown in Figure 5A. Alternatively, one could also prepare the data for a spatial analysis by having VALIS warp the cell positions from an existing dataset (Figure 5D). Figure 5C shows that VALIS can be used co-register H&E and IF images, which could be used to find ROI in the IF images, based on annotated H&E images.

Registration performed on CyCIF images was highly accurate, with an alignment error less than 10μm, which is about 1 cell diameter (Figure 4 and 5A). In these cases, the registration is accurate enough that cell segmentation and phenotyping could be performed. An example of such an analysis can be found in Figure 6A-C. HALO was used for cell segmentation and marker thresholding using the 32-channel image created by merging 11 rounds of registered CyCIF images (Figure 5A and 6A). For a full description of the channels, refer to Supplementary Table S1. A spatial analysis of the distribution of immune cells within the carcinoma region was conducted using 13 of the 32 markers, which were used to classify cells into one of nine cell types: helper T cells, cytotoxic T cells, regulatory T cells (Treg), natural killer (NK) T cells, active cytotoxic T cells (active CTL), memory T cells, M1 macrophages, M2 macrophages, B-cells, and tumor cells (Figure 6B, Table S2).

**Figure 5.**
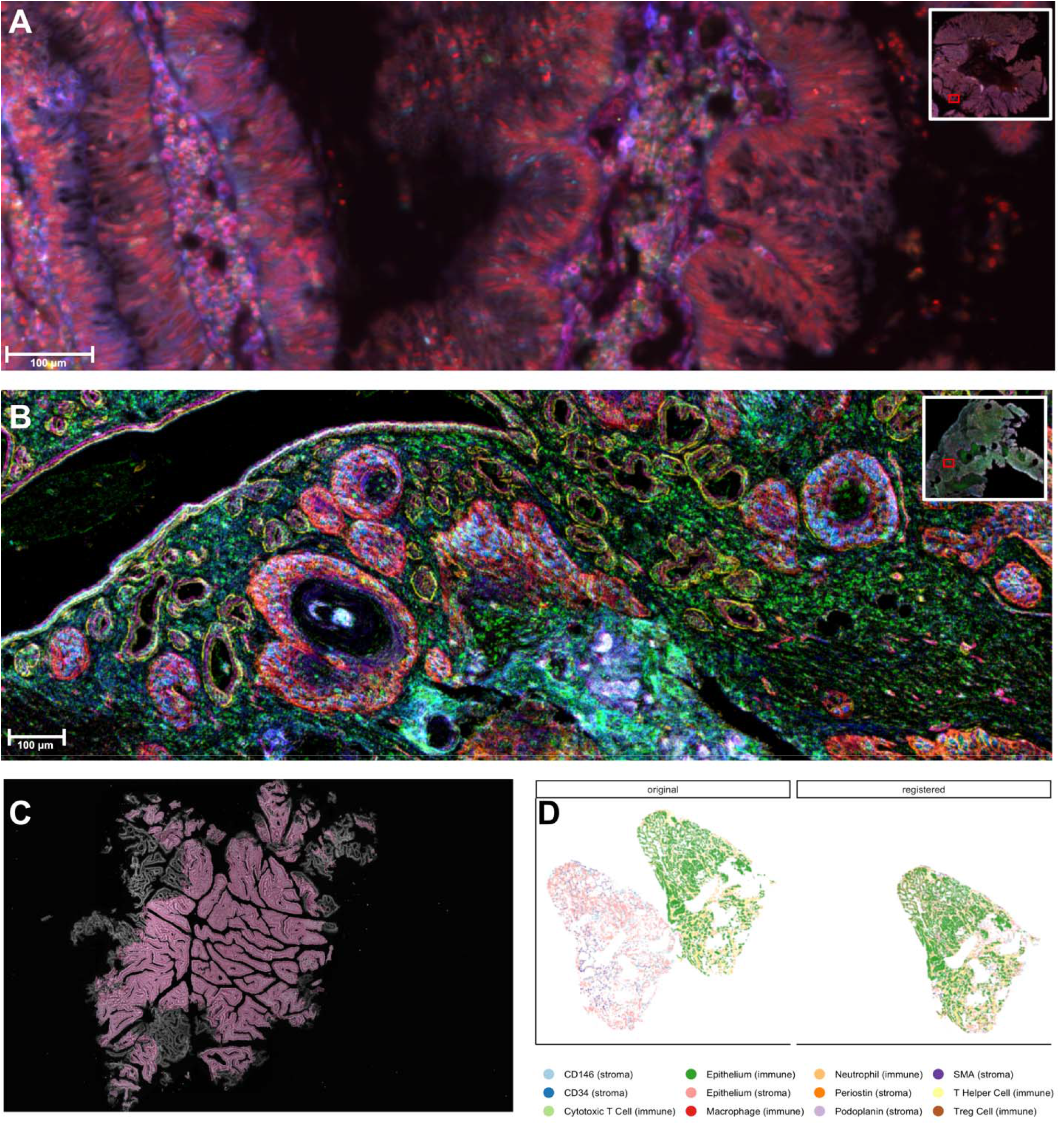
Potential applications of VALIS. **A**) Merging registered CyCIF slides, in this case creating a 32-channel image. **B**) Merging registered and processed IHC slides. Here, VALIS found the transformation parameters using the original images, but applied the transformations to 18 stain segmented versions slides (see Table S3 for list of markers). **C**) Registering an H&E slide with the DAPI channel of an IF slide, which may useful in cases where annotations H&E images would like to be used with IF images. Here, to visualize the alignment, the registered H&E image is overlaid on the DAPI channel. **D**) Applying transformations to cell segmentation data.

**Figure 6.**
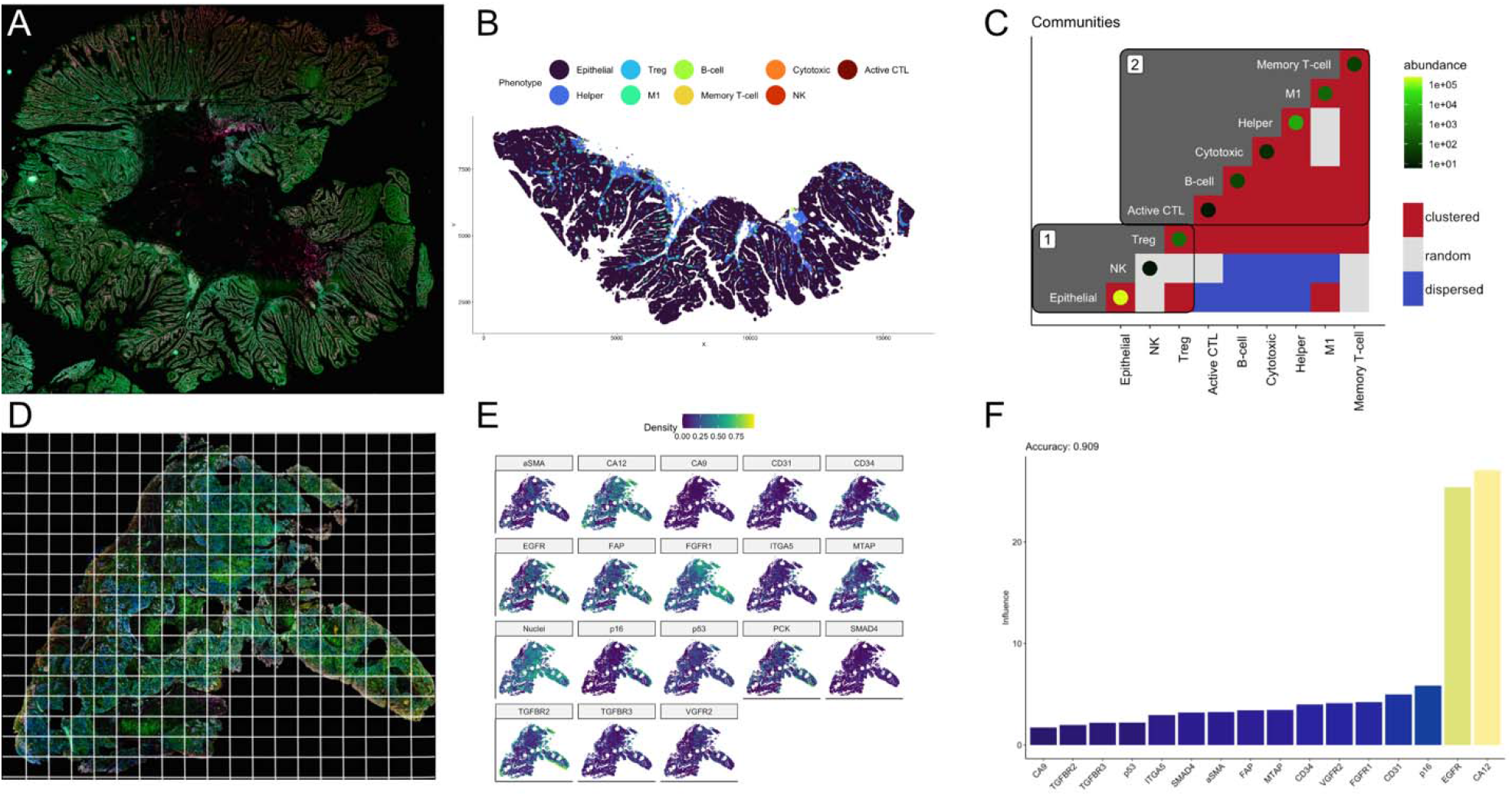
Example analyses using registered CyCIF WSI. A) A 32-channel image was created by registering and merging several rounds of CyCIF. The HALO platform was then used to perform cell segmentation and marker thresholding. B) Within the carcinoma region, a spatial analysis was conducted to determine the spatial relationship between 10 cell types, defined by different combinations of 13 markers. The pattern was determined using the DCLF test, where cell types could be found closer than expected (clustered), randomly distributed, or further apart than expected (dispersed). C) The observed patterns were used to construct a weighted network (1=clustered, 0=random, -1=dispersed), which subsequently underwent community detection. These results indicate the carcinoma (Epithelial) is largely isolated from the immune system. D) A composite IHC image of HNSCC using 18 markers of the tumor microenvironment. Alignment of IHC may not be cell-cell perfect, but using ecological methods, a spatial analysis can be conducted using quadrat counts. Each aligned slide underwent stain segmentation, the results of which were merged into a single composite image that was divided into regular quadrats. E) The number of positive pixels of each marker was calculated for each quadrat. F) A species distribution model was fit to the data to determine the role of each marker in creating a pro-tumor environment. Here, CA12 and EGFR were found to play the largest roles in creating a tumor supporting habitat.

A spatial analysis of the immune composition was conducted by first determining the spatial pattern observed between each pair of cell types (e.g. clustered, complete spatial randomness (CSR), or dispersion). Significance of departure from CSR was determined using the Diggle-Cressie-Loosmore-Ford (DCLF) test on the cross-type L-function for homogeneous point patterns (i.e. Besag’s transformation of Ripley’s K function) (Besag, 1977; Diggle, 1986; B. D. Ripley, 1977; B.D Ripley, 1981). These tests were conducted using the *spatstat* package for R (Baddeley, Rubak, & Turner, 2015; R Core Team, 2019). Clustering was considered significant when *p* ≤0.05 for the alternative hypothesis of “greater”, i.e. there were more cells within a radius *r* than expected under CSR. The spatial pattern was classified as dispersion when *p* ≤0.05 for alternative hypothesis of “lesser”. These patterns were then used to construct a weighted adjacency matrix, where 1=clustered, 0=CSR, and -1=dispersed (Figure 6C). The matrix was then divided into communities using the Leiden community detection algorithm (Traag, Waltman, & van Eck, 2019). This analysis revealed that the tumor (in community 1) is largely isolated from immune system (community 2).

Spatial analyses can also be conducted when alignments are not close enough for cell segmentation. One approach is to first divide the image into quadrats, and then count cells and/or quantify the markers in each quadrat. One can then can select from a wide variety of methods to conduct a spatial analysis of the quadrat counts. For example, one can create spatial association networks, species distribution models, and test for complete spatial randomness (Baddeley et al., 2015; Hijmans, Phillips, Leathwick, & Elith, 2017; Popovic, Warton, Thomson, Hui, & Moles, 2019).

Examples of spatial analyses of histological data with ecological methods based on quadrat counts or multiple subregions can be found in (C. Gatenbee et al., 2021; Chandler D. Gatenbee et al., 2019; C. D. Gatenbee, Minor, Slebos, Chung, & Anderson, 2020; Hunter, Moncada, Weiss, Yanai, & White, 2021; Maley, Koelble, Natrajan, Aktipis, & Yuan, 2015). Here, we provide a brief example using a sample that went through 18 stain/wash cycles, each time being stained for one of 18 tumor microenvironment (TME) markers (Figure 5B, 6D-F) (EGFR, H&E, FAP, α-SMA, TGFBR2, p16, FGFR1, TGFBR3, PCK, VGFR2, MTAP, CD34, CA9, p53, SMAD4, ITGA5, CA12, CD31). Each image underwent stain segmentation, the results of which were merged to create a single 18-channel composite slide (Figures 5B and 6D). This slide was then divided into 100μm x 100μm quadrats, and the number of positive pixels per quadrat for each marker was recorded (Figure 6A&B). A species distribution model was then fit to the quadrat counts, which allowed us to quantify the importance of each marker in creating a hospitable tumor microenvironment (Figure 6C). The results from this analysis indicate that EGFR and CA12 play the largest role in creating a pro-tumor microenvironment.

## Discussion

Here, we have provided a robust method to register IHC and/or IF WSI. Using this method, we have been able to increase the number of markers that can be included in spatial analyses. In the case of IHC, the maximum increase was from 1 marker to 69, and from 4 markers to 32 in CyCIF. Using the registered slides, we then provided examples of spatial analyses using the results yielded by VALIS, both when registration is close to cell-cell perfect, and when it is not.

VALIS can be used with existing datasets and protocols, as it works under a wide variety image resolutions and staining modalities, does not require H&E or DAPI stains, is non-destructive, does not require knowledge about the order slices were cut, and can be applied to both images and cell segmentation data. VALIS can therefore be used on tumor samples stored in archives that could not previously be analyzed by existing techniques. In addition to being applied to existing datasets, VALIS can also be used on datasets collected with spatial analysis in mind, such as CyCIF images. VALIS therefore provides a simple, fast, free, and opensource end-to-end solution to open, register, and save a wide variety of histology images that can subsequently undergo spatial analysis using a large number of cellular markers.

## Data availability

Images will be made available upon publication of the paper for which they were acquired.

## Code availability

Code will be made available on GitHub following publication.

## Acknowledgements

The authors gratefully acknowledge funding by the National Cancer Institute via the Cancer Systems Biology Consortium (CSBC) U01CA232382, the Physical Sciences Oncology Network (PSON) U54CA193489 and support from the Moffitt Center of Excellence for Evolutionary Therapy. The authors also wish to acknowledge the role of the Breast Cancer Now Tissue Bank in collecting and making available the samples used in the generation of this publication, and the patients who donated to the Bank.

## Supplemental

**Table S1.** Markers per registered CyCIF round. In addition to the markers lists, each round also had DAPI channel, which was used to register the rounds.

| CyCIF Round | Non-DAPI Channels |
| --- | --- |
| Round 1 | pHH3 |
|  | iNOS |
|  | CD45 |
| Round 2 | CD163 |
|  | CK |
|  | CD8 |
| Round 3 | CD44 |
|  | HLA-DR |
|  | PD-1 |
| Round 4 | Ecad |
|  | CD3 |
|  | CD20 |
| Round 5 | CD4 |
| | γ-H2AX |
|  | PD-L1 |
| Round 6 | CD45RO |
|  | FoxP3 |
| | α-SMA |
| Round 7 | p53 |
|  | CD68 |
|  | CD31 |
| Round 8 | CD11b |
|  | IDO1 |
|  | Vista |
| Round 9 | β-catenin |
|  | HLAABC |
|  | Myelo |
| Round 10 | CD45RB |
|  | Ki67 |
|  | CD57 |
| Round 11 | CD163 |
|  | CK |
|  | Vimentin |

**Table S2.** Cell phenotypes, and the makers used to define those phenotypes, used in the example spatial analysis using registered CyCIF rounds (Figures 5B, 6B-C).

| Phenotype | Marker(s) |
| --- | --- |
| helper T cells | CD3, CD4 |
| cytotoxic T cells | CD3, CD8 |
| regulatory T cells (Treg) | CD3, FOXP3 |
| natural killer (NK) T cells | CD3, DD57 |
| active cytotoxic T cells (active CTL) | CD3, CD8, HLA-DR |
| memory T cells | CD3, CD8, CD45RO |
| M1 macrophages | CD68, iNOS |
| M2 macrophages | CD68, CD163 |
| B-cells | CD20 |
| tumor cells | CK and/or E-cadherin |

**Table S3.** Markers used in example spatial analysis of registered IHC images (Figures 5B, 6D-F).

| Marker |
| --- |
| MTAP |
| CD34 |
| ITGA5 |
| p53 |
| p16 |
| FAP |
| EGFR |
| TGFBR2 |
| TGFBR3 |
| SMAD4 |
| FGFR1 |
| PanCK |
| CA9 |
| aSMA |
| VGFR2 |
| CD31 |
| CA12 |

